# Effects of IL2/anti-IL2 antibody complex on chikungunya virus-induced arthritis in a mouse model

**DOI:** 10.1101/2023.01.30.526329

**Authors:** Sarah R. Tritsch, Abigail J. Porzucek, Arnold M. Schwartz, Abigale M. Proctor, Richard Amdur, Patricia S. Latham, Gary L. Simon, Christopher N. Mores, Aileen Y. Chang

## Abstract

Chikungunya virus (CHIKV) is characterized by disabling joint pain that can cause persistent arthritis in approximately one-fourth of patients. Currently, no standard treatments are available for chronic CHIKV arthritis. Our preliminary data suggest that decreases in interleukin-2 (IL2) levels and regulatory T cell (Treg) function may play a role in CHIKV arthritis pathogenesis. Low-dose IL2-based therapies for autoimmune diseases have been shown to up-regulate Tregs, and complexing IL2 with anti-IL2 antibodies can prolong the half-life of IL2. A mouse model for post-CHIKV arthritis was used to test the effects of IL-2, an anti-IL2 monoclonal antibody (mAb), and the complex on tarsal joint inflammation, peripheral IL2 levels, Tregs, effector (Teff) T cells, and histological disease scoring. The complex treatment resulted in the highest levels of IL2 and Tregs, but also increased Teffs, and therefore did not significantly reduce inflammation or disease scores. However, the antibody group, which had moderately increased levels of IL2 and activated Tregs, resulted in a decreased average disease score. These results suggest the IL2/anti-IL2 complex stimulates both Tregs and Teffs in post-CHIKV arthritis, while the anti-IL2 mAb increases IL2 availability enough to shift the immune environment towards a tolerogenic one.

## Introduction

Chikungunya virus (CHIKV) is an alphavirus spread by the *Aedes aegypti* mosquito that is characterized by disabling joint pain that can cause persistent arthritis in approximately one-fourth of patients. For most CHIKV infections, acute symptoms typically resolve within 7–10 days; however, multiple studies have reported patients experiencing persistent joint pain months or years after the initial infection. In a study involving 485 CHIKV patients from Colombia, 25% reported persistent joint pain 20 months after infection, and 13% reported pain 40 months after infection [1,2]. The mouse model for post-CHIKV arthritis involves footpad inoculation of wild-type immunocompetent C57BL/6 mice, which causes localized swelling and systemic infection. In a pilot study involving the post-CHIKV arthritis mouse model, we determined that CHIKV footpad infection results in histologic evidence of arthritis, synovitis, periostitis, and myositis that persist to 21 days post-infection (dpi) [3].

Currently, there are no standard treatments available for chronic CHIKV arthritis. Our preliminary data suggest that decreases in interleukin-2 (IL2) during the acute stage of infection and the resulting alteration of regulatory T cell (Treg) function may play a role in CHIKV arthritis pathogenesis [3,4]. Novel low-dose IL2-based therapies for autoimmune diseases have been shown to up-regulate Tregs and may be of use in CHIKV arthritis flares in humans [5,6]. The short half-life of IL2 can be prolonged *in vivo* using various anti-IL2 monoclonal antibodies to form an IL2/anti-IL2 antibody complex [5,7]. Unlike other IL2 antibodies, the JES6-1 neutralizing antibody complexed with IL2 selectively induces Tregs expansion due to their high constitutive expression of the high-affinity IL2 receptor, IL2Rα (CD25). The specificity stems from the JES6-1 antibody binding to IL2 in a way that sterically hinders the interaction between IL2 and the low-affinity receptors IL2Rβ (CD122) and IL2Rγ (CD132) found on other immune cells [7,8].

We hypothesized that chronic CHIKV arthritis activity was associated with deficient IL2-mediated Treg levels; therefore, we tested the therapeutic effects of IL2, an anti-IL2 JES6-1 monoclonal antibody (mAb), and an IL2/anti-IL2 JES6-1 mAb complex on post-CHIKV arthritis in our mouse model. Our aims were 1) to describe the role of IL2 in Treg expansion and corresponding arthritis severity in CHIKV arthritis, and 2) to determine the role of IL2 therapy in treating CHIKV arthritis in a mouse model.

## Results

### Viremia and tarsal joint inflammation

At two days post-infection (dpi), all mice were confirmed CHIKV-positive by qRT-PCR. By 14 dpi, most mice were CHIKV-negative, and by 16 dpi, all mice were confirmed CHIKV-negative.

Tarsal joint measurements exhibited an overall increase in infected tarsal joints’ swelling that peaked at six dpi (Figure 2). While all treatment groups returned to baseline by the end of the study, the IL2-treated group showed the steepest drop in inflammation between 16 and 17 dpi. During IL2 treatment (days 16–19), there was a trend level time effect (p = 0.07) with a reduction over time, on average. However, the interaction between the treatment group and day was insignificant (p = 0.41), indicating that the change over time was similar between treatment groups. Results were similar using the inflammation measure, calculated as a change from baseline joint size divided by baseline joint size (Figure 2B).

**Figure 1.**
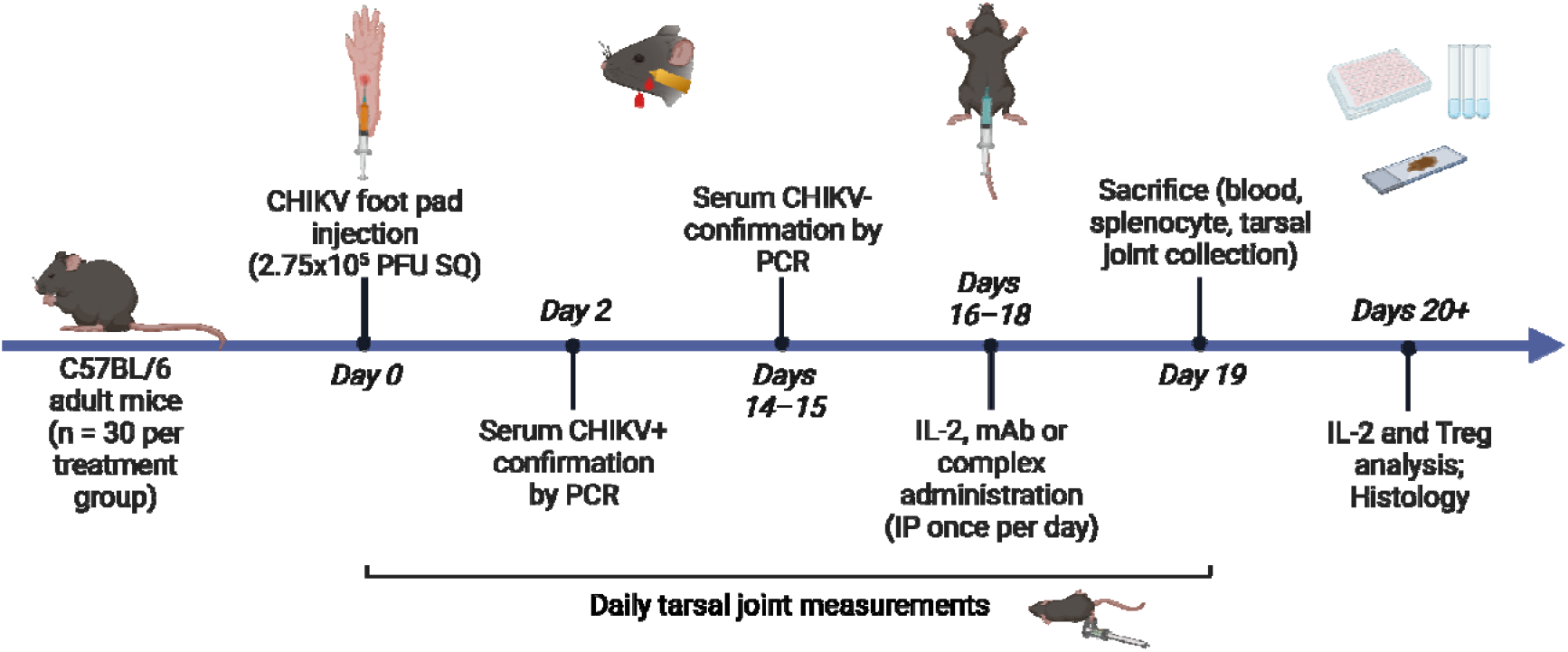
CHIKV post-acute arthritis mouse model with IL-2-based treatments. C57BL/6 adult mice were infected with CHIKV via footpad injection. Mice were confirmed positive for CHIKV at two days postinfection (dpi) and confirmed negative by 16 dpi. Recombinant IL-2, anti-IL-2 monoclonal antibody (mAb), or IL-2/anti-IL-2 mAb were administered via intraperitoneal injection once per day for three days, starting at 16 dpi. Mice were sacrificed at 19 dpi, and spleen, hind limbs, and blood were collected for future analysis. Tarsal joints were measured daily.

**Figure 2.**
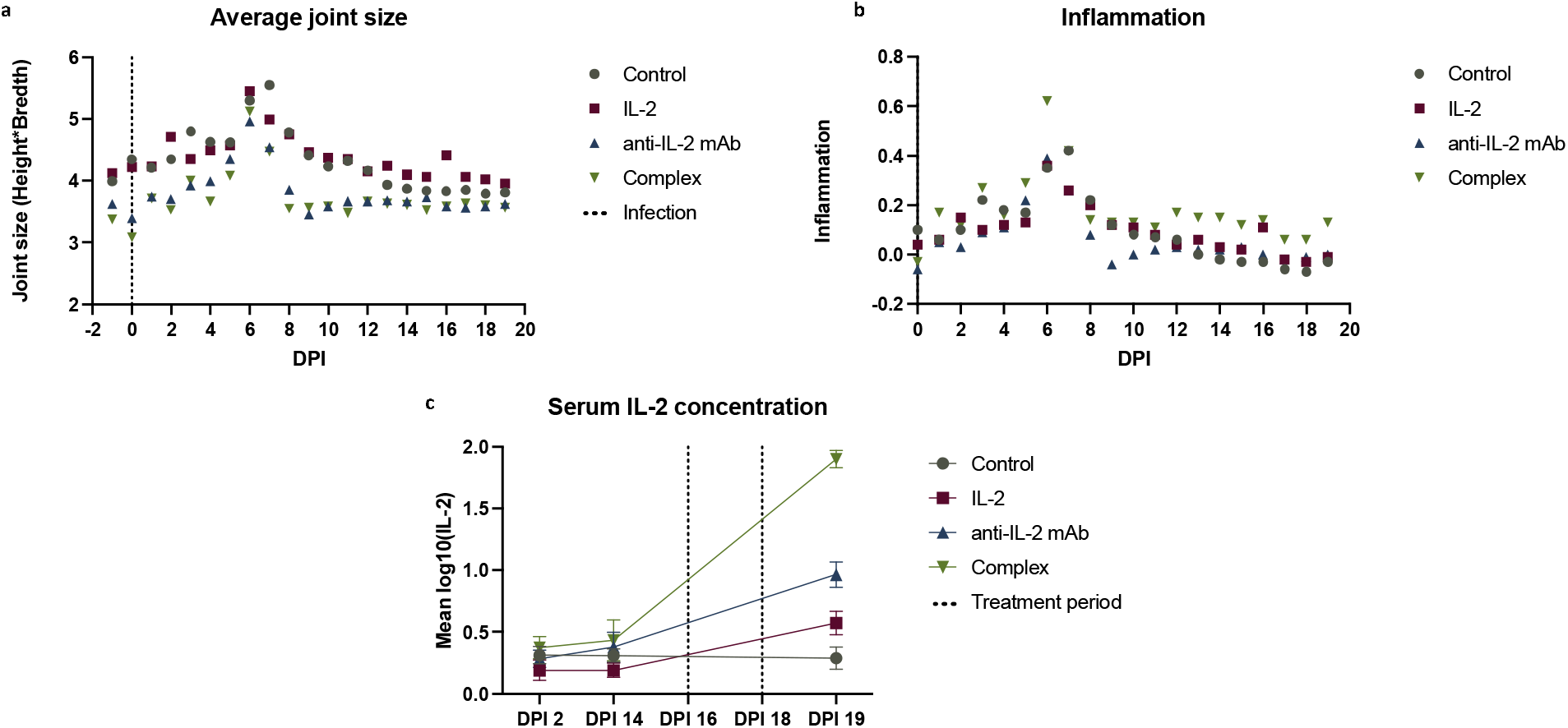
Tarsal joint measurements, inflammation, and serum IL2 levels. A) Average joint size of mice by treatment group on each day of the study, starting one day before infection, which served as the baseline joint size for calculating inflammation. The average joint size was calculated by multiplying the height by the breadth of the left tarsal joint. B) Tarsal joint inflammation was calculated using [(x-day 0)/day 0)], where x is the footpad size measurement for a given day post-infection. C) Serum IL-2 concentrations were normalized using log10(IL-2 + 1). Vertical lines illustrate the treatment period.

### IL2 levels

IL2 distributions were normalized using log10(IL2 + 1) for analysis. IL2 levels were low and stable in all treatment groups between 2 and 14 dpi (before treatment). However, at 19 dpi (after treatment), IL2 levels increased significantly compared to 14 dpi (before treatment) in the three experimental groups, with a 2.3-fold increase in IL2-treated mice, a 6.9-fold increase in IL2 mAb-treated mice, and a 57.8-fold increase in complex-treated mice compared to the control group (Figure 3). IL2 levels in the control group did not change. In the mixed model examining change in IL2 level from day 14 to 19 by treatment group, the main effects of the treatment group and time were both significant (both p < 0.0001), as was the interaction between treatment group and time (p < 0.0001).

**Figure 3.**
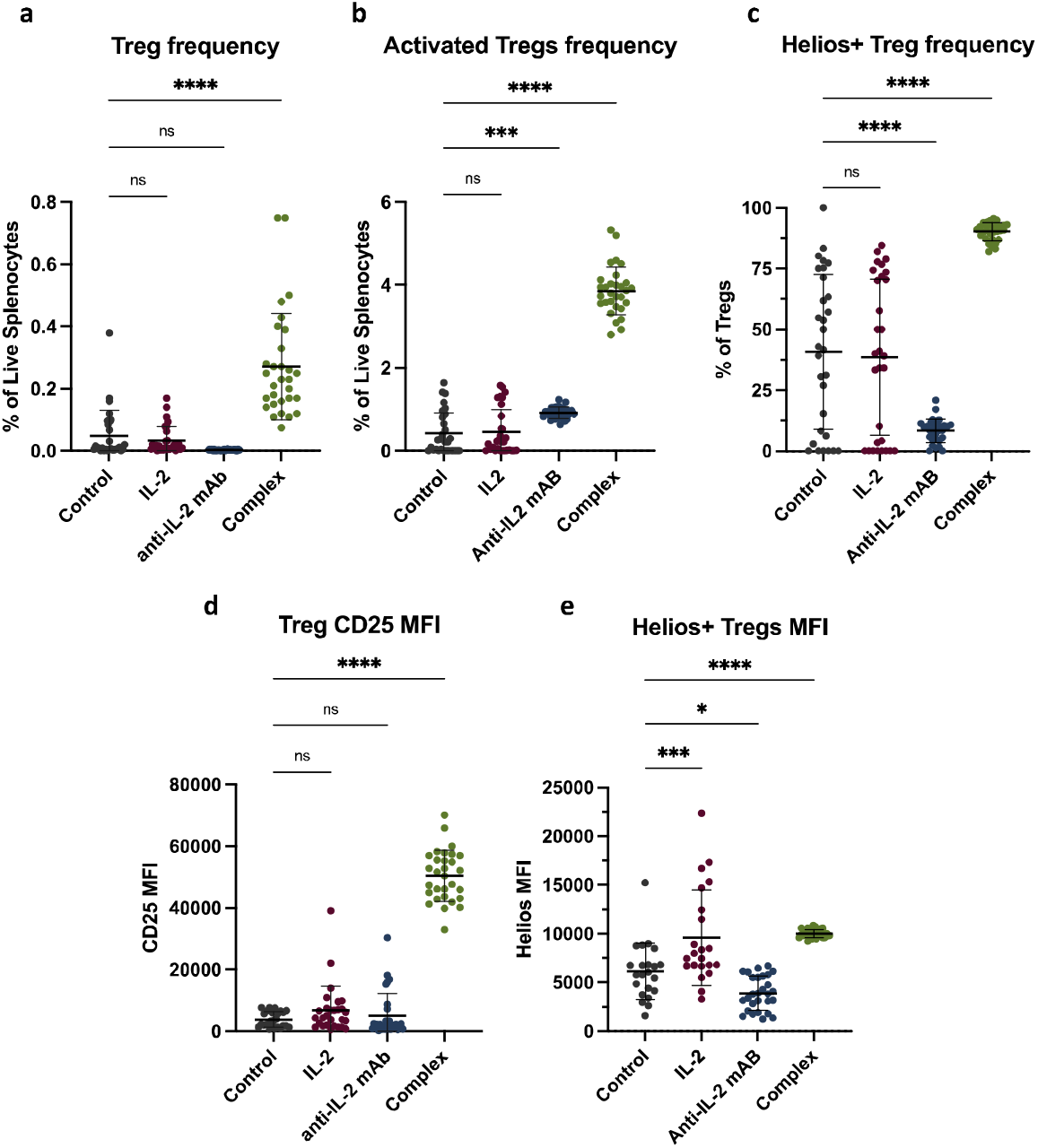
Regulatory T cell flow cytometry performed on splenocytes. A) CD4+CD25^hi^FoxP3+ Treg cell frequency out of live splenocytes. B) FR4^hi^CD25^hi^ activated Treg cell frequency out of live splenocytes. C) Frequency of Helios-positive Tregs out of all Tregs. D) CD25 median fluorescence intensity (MFI) on Tregs. E) Helios MFI on Helios+ Tregs.

### Treg proliferation and activation

Table 1 summarizes the flow cytometry performed on splenocytes, which determined that Treg frequency and activation levels differed significantly across all treatment groups (p < 0.0001, Figure 3). There was a significant increase in the percent Tregs out of live splenocytes in complex-treated mice compared to controls (p < 0.0001). When FR4 and CD25 were used to determine the level of activated Tregs, a slight increase in antibody-treated mice and a greater increase in complex-treated mice was observed compared to the control group (p < 0.0001). There was also a significant increase in the Helios activation marker in complex-treated mice and a significant decrease in antibody-treated mice compared to the control group (p < 0.0001).

**Table 1.**
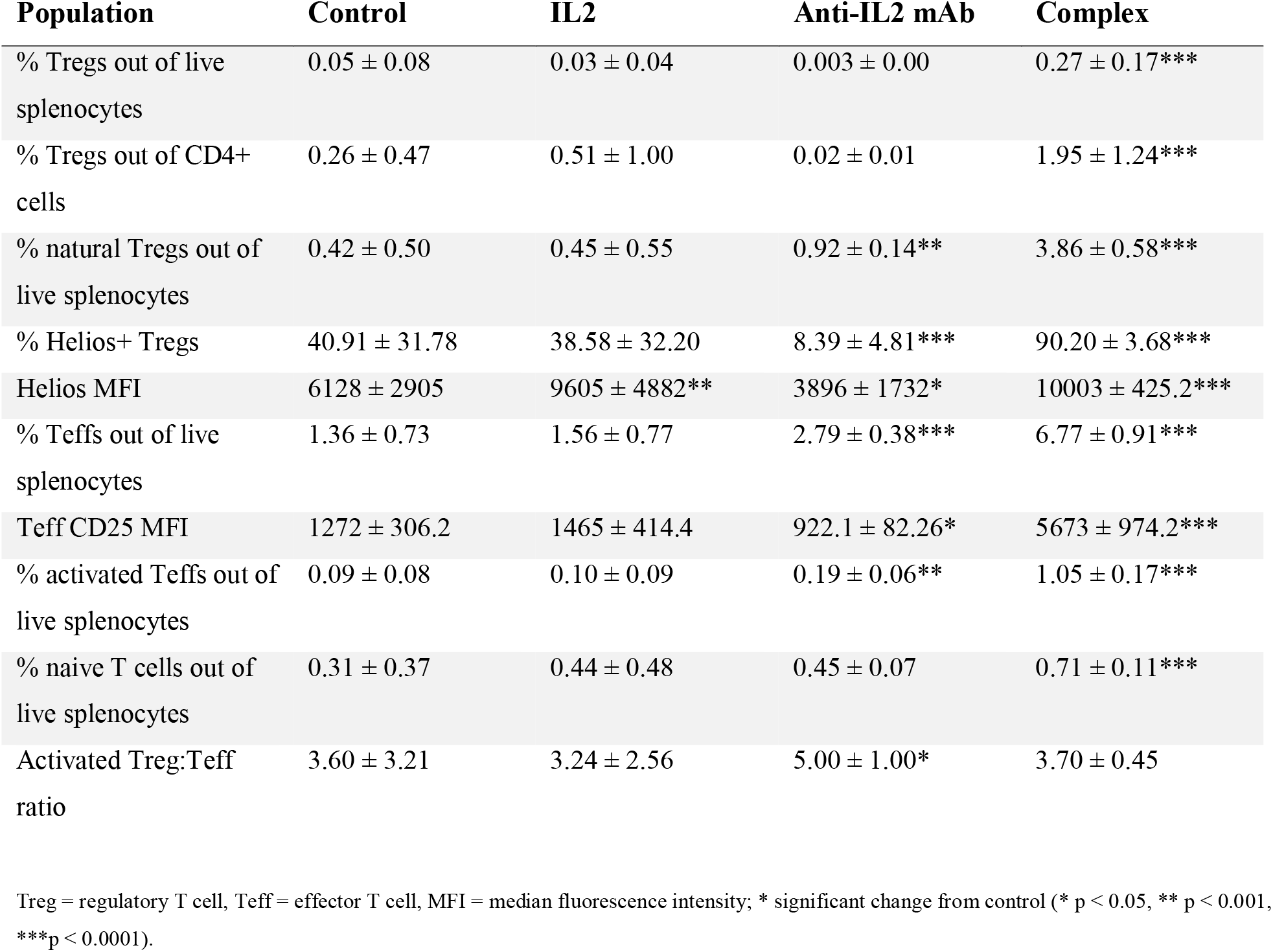
Regulatory and effector T cell levels by treatment group (mean ± SD) determined by flow cytometry on mouse splenocytes.

| Population | Control | IL2 | Anti-IL2 mAb | Complex |
| --- | --- | --- | --- | --- |
| % Tregs out of live splenocytes | 0.05 ± 0.08 | 0.03 ± 0.04 | 0.003 ± 0.00 | 0.27 ± 0.17*** |
| % Tregs out of CD4+ cells | 0.26 ± 0.47 | 0.51 ± 1.00 | 0.02 ± 0.01 | 1.95 ± 1.24*** |
| % natural Tregs out of live splenocytes | 0.42 ± 0.50 | 0.45 ± 0.55 | 0.92 ± 0.14** | 3.86 ± 0.58*** |
| % Helios+ Tregs | 40.91 ± 31.78 | 38.58 ± 32.20 | 8.39 ± 4.81*** | 90.20 ± 3.68*** |
| Helios MFI | 6128 ± 2905 | 9605 ± 4882** | 3896 ± 1732* | 10003 ± 425.2*** |
| % Teffs out of live splenocytes | 1.36 ± 0.73 | 1.56 ± 0.77 | 2.79 ± 0.38*** | 6.77 ± 0.91*** |
| Teff CD25 MFI | 1272 ± 306.2 | 1465 ± 414.4 | 922.1 ± 82.26* | 5673 ± 974.2*** |
| % activated Teffs out of live splenocytes | 0.09 ± 0.08 | 0.10 ± 0.09 | 0.19 ± 0.06** | 1.05 ± 0.17*** |
| % naive T cells out of live splenocytes | 0.31 ± 0.37 | 0.44 ± 0.48 | 0.45 ± 0.07 | 0.71 ± 0.11*** |
| Activated Treg:Teff ratio | 3.60 ± 3.21 | 3.24 ± 2.56 | 5.00 ± 1.00* | 3.70 ± 0.45 |

Treg = regulatory T cell, Teff = effector T cell, MFI = median fluorescence intensity; \* significant change from control (\* $p < 0.05$ , \*\* $p < 0.001$ , \*\*\* $p < 0.0001$ ).

Teff cell frequency (Figure 4) increased significantly in both the complex-treated and the antibody-treated groups compared to the control group (p < 0.0001). The CD25 median fluorescence intensity (MFI) on the Teff cells decreased slightly with marginal significance in the antibody group and increased more than 4-fold in the complex group compared to the control group (p < 0.0001). Additionally, there was a slight increase in the number of activated Teff in the antibody group and a more considerable increase in the complex group compared to the control group (p < 0.0001), as well as a significant rise in naïve T cells in the complex group compared to the control group (p < 0.0001). Finally, there was a moderate increase in the activated Treg to activated Teff ratio in the antibody group alone. There were no significant changes in the frequency of Tregs, Helios-positive Tregs, Teff, or CD25 MFI of Teffs in the IL2 group compared to the control group.

**Figure 4.**
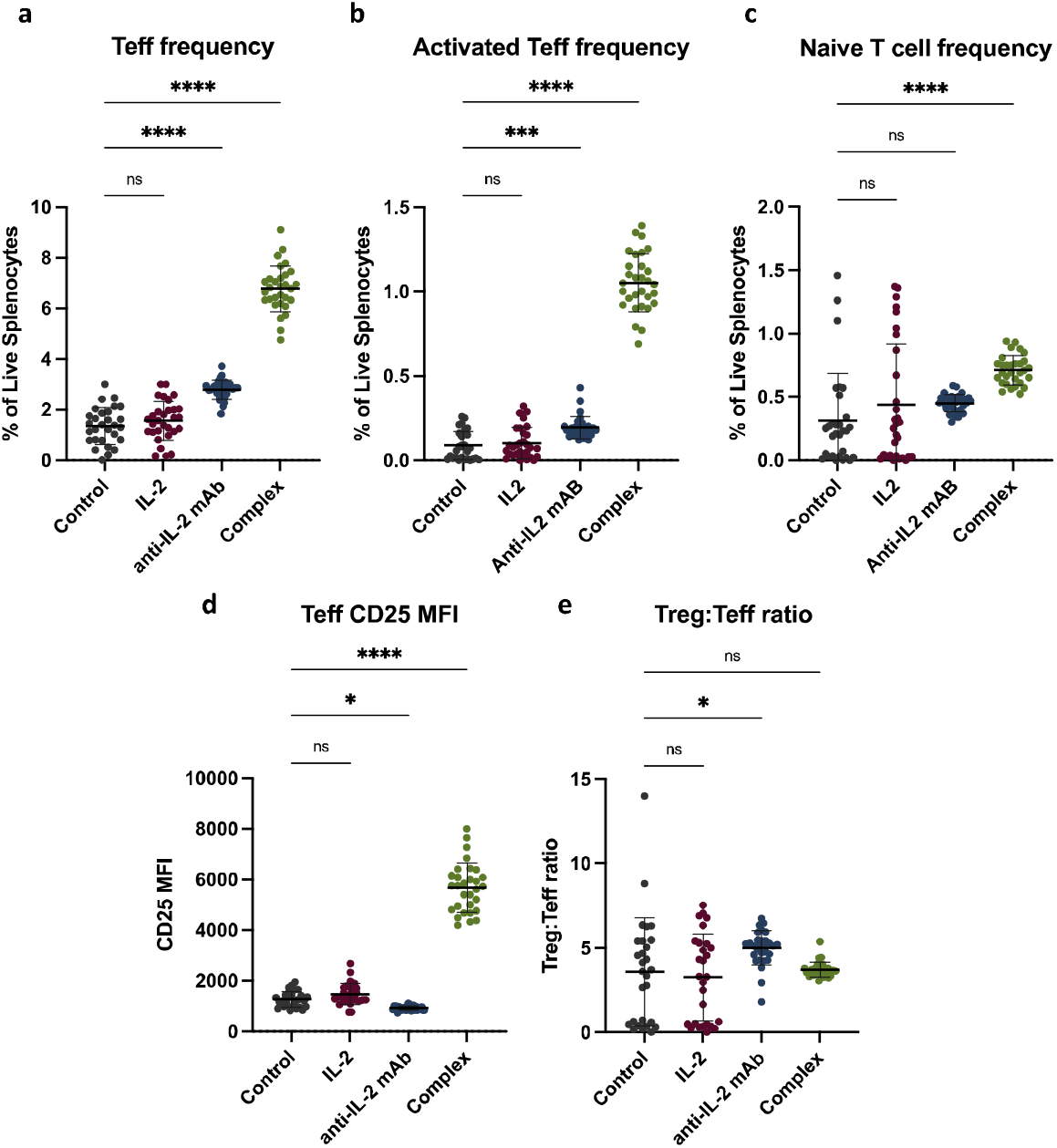
Effector and naïve T cell flow cytometry performed on splenocytes. A) Frequency of CD4+CD25+FoxP3-Teff cells out of live splenocytes. B) FR4^int^CD25^int/hi^ activated Teff cell frequency out of live splenocytes. C) FR4^lo^CD25^lo^ naïve T cell frequency out of live splenocytes. D) CD25 median fluorescence intensity (MFI) on Teffs. E) Ratio of activated Treg cells to activated Teff cells.

### Histology

For each mouse, two independent ratings were done for the individual and composite histologic scoring. Three methods were used to evaluate the test-retest reliability. First, the Pearson correlation coefficient (r) for reliability between ratings was examined with an r > 0.90 set as acceptable. The obtained r-value was 0.904, indicating good reliability. Next, the Bland-Altman plot was reviewed to determine the fixed bias, which ideally should be zero. The fixed bias was calculated as 0.06, indicating good reliability. Finally, the weighted kappa was examined as a measure of inter-rater agreement. The weighted kappa was 0.78, indicating acceptable reliability (95% confidence interval 0.71–0.85).

Figure 5 displays representative images of healthy (A) and CHIKV-infected (B–D) tarsal joints. Composite histology scores of left (infected) tarsal joints revealed a higher number of mice with severe arthritis in control (4 mice) and IL2 (6 mice) groups compared to the antibody (0 mice) and complex (1 mouse) groups (Figure 5E). Additionally, after adjusting for sex, the mean composite histology score was significantly lower in the antibody group than in any other group (p = 0.03). The antibody group was the only group with composite histology scores significantly different from the control group (p = 0.0067). When looking at individual components of the histology, mice in all groups presented with synovitis and a majority had at least minor myositis and periosteal inflammation, while few mice showed cartilage or bone erosion (Supp. Fig. 1).

**Figure 5.**
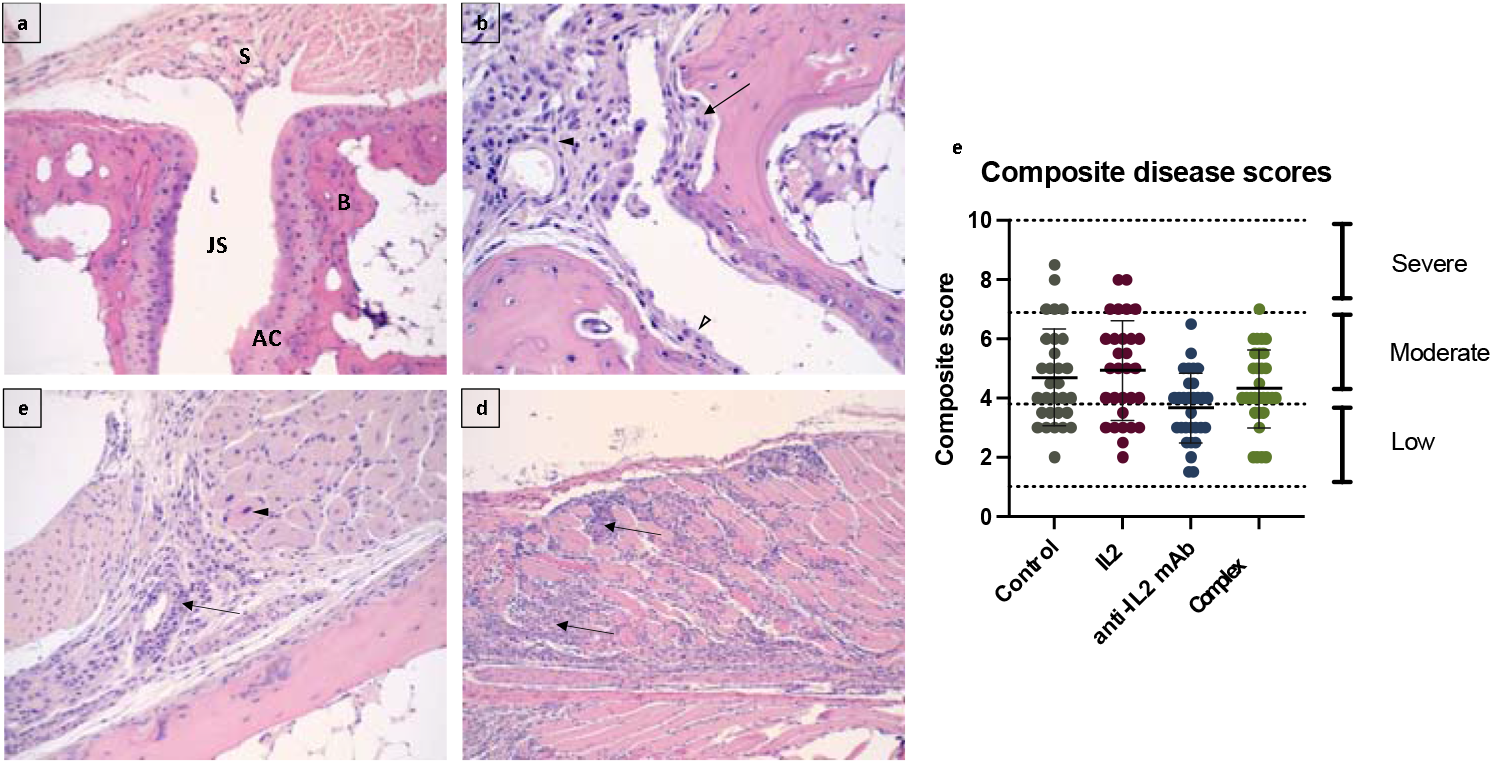
Histology images of mouse tarsal joints. A) Healthy tarsal joint illustrating the synovium (S), joint space (JS), articular cartilage (AC), and subchondral bone (B); B) CHIKV+ tarsal joint with bone and articular cartilage erosion (arrow), active synovitis (solid arrowhead), and pannus with articular cartilage erosion (open arrowhead); C) CHIKV+ tarsal joint with mild myositis with degenerative myocytes (arrowhead), fasciitis (arrow), and periostitis with cortical bone erosion; and D) CHIKV+ tarsal joint with severe myositis with degeneration and necrosis (arrows). E) Composite histological disease scores for each mouse by treatment group. Scores for each mouse were calculated by adding together the scores from each histological component. A score of 0 to <4 was considered low severity, 4 to <7 was moderate, and >7 was severe.

### Mouse sex effects on outcomes

The effects of mouse gender on the data were investigated. While males had larger joint sizes than females at baseline and over all time points, the sex effect on inflammation during treatment (days 16–19) was not significant (p = 0.14), indicating that overall inflammation was similar for male and female mice. The sex-by-day interaction was insignificant (p = 0.65), indicating that the change in inflammation during treatment did not differ by sex. Further, none of the Treg levels were significantly associated with sex, and controlling for sex did not affect the association of the treatment group with Treg levels.

However, in the IL2 model comparing day 14 (before treatment) to day 19 (after treatment), the 3-way interaction of the treatment group-by-time-by-sex was significant (p < 0.0001), indicating that the group-by-time interaction differed by sex. While the change in peripheral IL2 levels over time in control and IL2 treatment groups is similar for males and females, females had a more considerable increase in IL2 over time, but only in the antibody and complex treatment groups. Additionally, the composite score was significantly higher for females after controlling for the treatment group (p < 0.0001), and the group effect remained significant after controlling for sex (p = 0.01). Proportions of mice with low, medium and high severity were 80%, 17%, and 3%, respectively, for males versus 42%, 43%, and 15% for female mice (p < 0.0001).

## Discussion

The investigation into the use of IL2-based therapeutics to treat post-CHIKV arthritis yielded interesting results. Daily tarsal joint measurements lacked significant changes between groups, which was expected based on our previous study [3]. IL2 analysis indicated stable levels of peripheral IL2 before treatment across all groups. After treatment, there was a small but significant increase in IL2 concentrations in the IL2 group, a larger increase in the antibody group, and the most substantial increase in the complex group. Interestingly, the antibody group was the only one with a significant decrease in the average composite disease severity score. Conversely, flow cytometry analysis of splenocytes revealed that the complex was the only treatment to significantly increase the frequency and activity of FoxP3+ Tregs. When populations of activated Tregs defined as FR4^hi^CD25^hi^ were considered, there was a small but significant increase in the antibody group and a large increase in the complex group. There was a significant increase in the Teff cells in both the antibody and complex groups, but a decrease in Teff cell CD25 expression in the antibody group and an increase in the complex group, which is a marker of Teff activation. Consequently, the antibody group was the only group with a higher activated Treg to activated Teff cell ratio, pointing to a more tolerogenic immune environment compared to the other groups. Finally, sex analysis of the data revealed that females had worse histology outcomes, corresponding to human disease; however, females in the antibody and complex groups were more responsive to IL2 than males. This could be due to differences in size since male mice are typically larger than female mice.

Figure 6 depicts IL-2 and T-reg immunology in normal function and active CHIKV arthritis. The left panel illustrates the interaction of normal peripheral levels of IL-2 and Tregs during normal suppressive function. In homeostasis, IL-2 binds to the high-affinity receptor (CD25) on Tregs, which drives their expansion and activation. Tregs can then suppress the growth of effector T cells, thereby controlling inflammation. As shown in the middle panel, deficient IL-2 levels in CHIKV arthritis lead to a lack of Treg expansion and activation, allowing T effector cells to expand unregulated and leading to increased local inflammation [4,9]. We hypothesized that IL-2-based treatments such as recombinant IL-2, an anti-IL-2 mAb, or the two in complex would increase peripheral levels of IL-2, therefore increasing the proliferation and activation of Tregs and depressing Teff cells.

**Figure 6.**
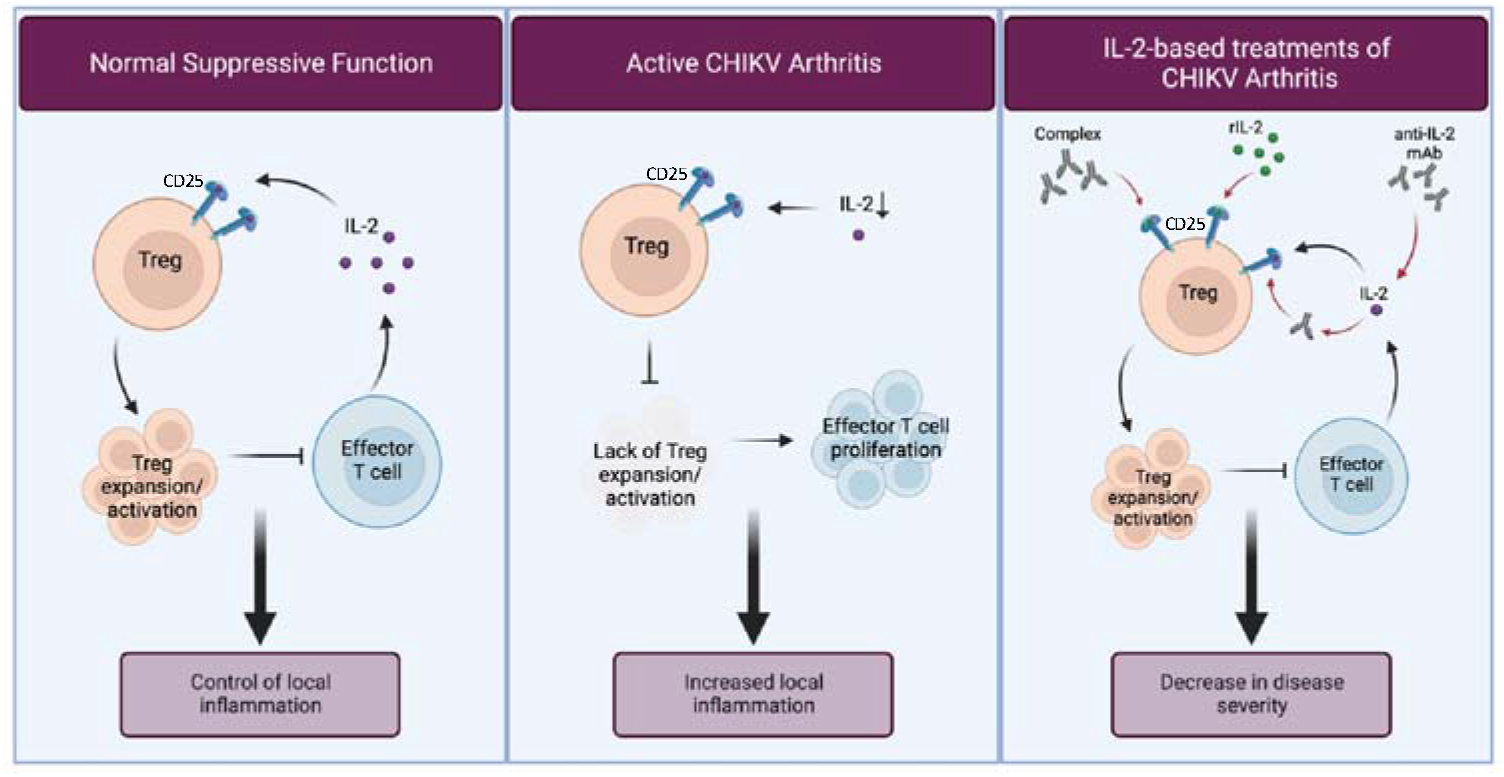
IL-2 and T-reg immunology in normal immune function and active CHIKV arthritis.

Previous studies by Lee et al. (2015 and 2016) have investigated the effects of the IL2/anti-IL2 complex administered before and at the peak of CHIKV infection in mice. Prophylactic treatment with the complex prevented arthralgia development, while treatment during acute infection caused infiltration of pathogenic CD4+CD25+ Teff cells and intensified joint inflammation [6,10]. Our study aimed to determine the complex’s effects on active arthralgia after viral clearance, as this is when patients usually present for treatment of post-CHIKV arthritis. However, treatments had to be administered immediately after the virus was cleared since, unlike in humans, diseased joint pathology in mice will eventually clear on its own. Based on the results from Lee et al., it is possible that pathogen-activated Teff cells, which express high levels of CD25, were still in circulation in our mice, even after the virus was cleared. Indeed, Teff populations in the complex group had significantly higher CD25 expression, while the antibody group Teff populations had significantly lower CD25 expression. Additionally, the frequency of activated Teff cells defined as FR4^int^CD25^int/hi^ moderately increased in the antibody group, with a 10-fold increase in the complex group.

The histologic examination of the lower limb provided insight into the nature and progression of the inflammatory process. Encouragingly, none of the treatments worsened the disease outcome. Nonetheless, all treatment groups had mice with low, moderate, and severe disease scores. This is to be expected, as variation in disease progression after infection can occur among individual mice. Histologic scoring also demonstrated that synovitis was an early and prominent process, given that no case (control or therapeutic) was free of synovitis. Moreover, when synovitis was mild (individual histologic score of 1), there was no case of overall composite histologic severity (overall histologic score of 7, 8, 9, or 10). Similarly, when the synovitis score was marked (individual histologic score of 2), three-quarters of the cases had overall histologic scores of five or greater, indicative of moderate to severe disease. Further, most mice had at least mild myositis and periosteal inflammation. Few mice, however, showed signs of cartilage or bone erosion. This suggests that the disease pathology in mice begins with synovitis, progressing to muscle and periosteal involvement, and finally, cartilage and bone erosion in cases of severe disease. The variability of disease expression indicates that disease progression and host response pathways for injury and inflammatory reactivity differ in their extent among the cohort of cases.

The data suggest that complexing the IL2 with the anti-IL2 mAb makes it more biologically available and thus, has a more significant impact on the IL2-mediated increase in Tregs, which decreases overall inflammation and joint degradation. However, there seems to be a goldilocks effect associated with these treatments, in which low-dose IL2 alone is insufficient to drive a robust Treg response, while the complex increases IL2 levels too much, affecting the frequency and activity of Teff cells in addition to Tregs. This could be because the antibody concentration was not high enough to bind all the available exogenous IL2 in the complex or because it allowed endogenous IL2 the opportunity to bind low-affinity receptors on Teff cells. The IL2 antibody, on the other hand, increased IL2 levels just enough to act on Treg cells but not Teff cells. We hypothesize that since the JES6-1 anti-IL2 antibody recognizes the IL2Rβγ (CD122) site of IL2, administration of the antibody alone binds available endogenous IL2, extending its half-life and ensuring its delivery to Treg cells [11]. Conversely, when JES6-1 is complexed with IL2, endogenous (or excess exogenous) IL2 is free to bind low-affinity receptors on Teff cells, driving their proliferation and activation.

A few limitations were noted during this study. First, treatments were not weight-based, which could account for the differences in response to therapies between male and female mice. Additionally, this study used a post-acute mouse model, which differs from the clinical progression of post-acute human disease. A dose-response study in this mouse model could elucidate the anti-IL2 antibody mechanism for decreasing post-acute disease. Further, co-administration of the complex with a Teff cell inhibitor, such as rapamycin, could dampen the nontarget effects of the primary treatment [12]. Ultimately, these highly impactful findings indicate the need for human *in vitro* and clinical studies evaluating anti-IL2 monoclonal antibodies and IL2/anti-IL2 complex to treat CHIKV-related chronic arthritis.

## Methods

### Ethics statement

All animal care and procedures performed in this study were reviewed and approved by the George Washington University (GWU) Institutional Animal Care and Use Committee (IACUC) protocol #A456, which complies with the guidelines stated in the National Institutes of Health’s (NIH) Guide for the Care and Use of Laboratory Animals. This manuscript was written in accordance with the ARRIVE guidelines (Animal Research: Reporting of In Vivo Experiments)

### Mice

A total of 120 mice were included in this study. Equal numbers of 10-week-old male and female C57BL/6J mice were obtained from The Jackson Laboratory (https://www.jax.org/) and housed in groups of five in micro-isolator cages in the ABSL-3 laboratory at GWU. Mice were given free access to food and water.

Mice were anesthetized with isoflurane, then injected with 25 μl (2.75 × 10^5^ plaque forming units (PFU)) of CHIKV subcutaneously in the caudoventral aspect of the left hind foot near the tarsal joint. Age and sex-matched negative control mice were injected with 25 μl of sterile phosphate buffered saline (PBS). Mice were monitored at least once daily for signs of illness, such as lethargy, joint swelling, lameness, lethargy, and limb gnawing. At 19 dpi, all mice were euthanized via CO_2_ asphyxiation, followed by a cardiac stick. Figure 1 illustrates the stepwise flow of the protocol.

### Virus

The strain of CHIKV used for this study was from the East, Central, and South African (ECSA) lineage that was initially isolated from a human case in South Africa (SAH-2123) and was supplied by the World Reference Center for Emerging Viruses and Arboviruses (Galveston, TX, USA). This strain was used in a previous study performed by our lab to determine a mouse model for studying chronic CHIKV arthritis [3].

All studies performed with viable CHIKV were performed in Biosafety Cabinets in a certified BSL-3 laboratory and were conducted with the approval of the GWU Institutional Biosafety Committee protocol #IBC-19-026.

CHIKV was passaged once in Vero cells (CCL-81, ATCC) in DMEM containing antibiotics and antimycotics, then quantified by viral plaque assay and quantitative real-time – polymerase chain reaction (qRT-PCR). The calculated titer of the stock virus was 1.1E7 PFU/ml, yielding a predicted limit of detection for the qRT-PCR assay of 0.01 PFU/ml.

### Viremia detection

At 48 hours post-infection, 200 μl of blood was collected from the submandibular vein for serum analysis to confirm mice confirmed CHIKV positive by qRT-PCR. The Zymo Research Quick DNA/RNA Kit was used to isolate viral RNA, and the viral load was quantified using the Invitrogen Superscript III Platinum One-Step qRT-PCR Kit and CHIKV primer/probe set [13]. RT-PCR was analyzed using a LightCycler 96 Instrument (Roche Diagnostics) with thermal cycling conditions as follows: one cycle at 50°C for 900 sec, one cycle of 95°C for 120 sec, and 45 cycles of 95°C for 3 sec, followed by 55°C for 30 sec and a cooling cycle of 37°C for 30 sec.

### Treatment with IL2 and monoclonal antibody

At 16 dpi (the point at which all mice were CHIKV-negative), mice were treated once per day for three days by intraperitoneal injection with phosphate buffered saline (PBS) as a control, 1.5 μg/day recombinant mouse IL2, 5 μg/day anti-IL2 monoclonal antibody (mAb; clone JES6-1), or an IL2/anti-IL2 complex containing 1.5 μg of IL2 and 5 μg IL2 mAb. Thirty mice were included in each treatment group (15 males and 15 females).

### Serum IL2 analysis

Peripheral blood was collected from the submandibular vein at 2, 14, 16, and 18 dpi and via cardiac puncture at 19 dpi. Blood was allowed to clot at room temperature for 30 minutes, then centrifuged at 3,000 x g for 10 minutes. Serum was aliquoted and stored at −80°C until analysis. The IL2 Mouse ProQuantum Immunoassay Kit (Invitrogen) was used to analyze serum IL2 levels.

### Flow cytometry

Spleens were dissected from mice following euthanasia, and splenocytes were collected by mashing the spleens through a 70-um cell strainer. Red blood cells were lysed using AKT lysis buffer. Cells were then stained for flow cytometry using the Biolegend antibodies FITC CD4 (clone RM4-5), PE-Cy7 CD25 (clone PC61.5), PE FoxP3 (clone MF-14), APC Helios (clone 22F6), PerCP-Cy5.5 FR4 (clone 12A5), and Fixable Live/Dead Aqua stain from Invitrogen, as well as the FoxP3 Fixation/Permeabilization kit from eBiosciences. A BD LSRII flow cytometer was used to process samples, and analysis was performed using FlowJo Software. Treg cells were defined as CD4+CD25^hi^FoxP3+, and Teff cells were defined as CD4+CD25+FoxP3-. Further, FR4 (folate receptor 4) was used in combination with CD25 staining to define activated Tregs (FR4^hi^CD25^hi^), activated effector T cells (FR4^int^CD25^int-hi^), and naïve T cells (FR4^lo^CD25^lo^) [14].

### Histology staining

After euthanasia, hindlimbs were dissected and fixed in 10% formaldehyde at 4°C for a minimum of 48 h. Fixed tissues were rinsed with distilled water, and any remaining skin was removed. Decalcification of bones was achieved using a solution containing EDTA, sodium hydroxide, and polyvinylpyrrolidone (PVP) with a pH of 7.0–7.4. Tissues were submerged in the decalcification solution at 4°C with agitation for 28 days, and the solution was changed every other day to maintain a pH of 7.0–7.4.

After decalcification, tissues were rinsed with distilled water and processed in PBS three times for 5 min each, in distilled water three times for 5 min each, in 50% ethanol for 5 min, and finally, in 70% ethanol for 5 min. After the final wash, the tissues were stored in 70% ethanol at 4°C until processed.

To process for histopathological evaluation, tissue samples were dehydrated in increasing ethanol concentrations, xylene, and embedded in paraffin wax, from which five μm-thick sections were adhered to positively charged glass slides. Samples were stained with hematoxylin and eosin (H&E) for evaluation, and Masson’s trichrome stains were performed on select slides. All histologic evaluations were performed by a board-certified pathologist, who was blinded to the treatment group. Two blinded reads were performed one month apart, and the histologic score was averaged. All slides were evaluated using an Olympus BK46 microscope, and digital images were obtained using a Spot Imaging digital camera and Spot Pathsuite 2.0.

### Histologic inflammation scoring system

The generation of the semi-quantitative histologic inflammation scoring system used in this study has been previously described [3]. In short, a microscopic evaluation of bone and joint tissue assessed inflammatory reaction and injury in (1) the synovium, (2) articular cartilage, (3) skeletal muscle and soft tissue, (4) periosteum, and (5) cortical bone. The scoring ranged from zero (no injury/inflammation) to two (significant injury/inflammation) for each of the five components, with a total possible composite score of 0 to 10. The scoring system was based on the maximal degree of inflammation and injury seen on day seven in our pilot study [3].

### Tarsal joint measurement

Tarsal joint width and breadth were measured daily using calipers and recorded in mm. Each joint measurement was performed three consecutive times and averaged to account for variability. The width was measured directly posterior to the tarsal joint foot pad projection, with only enough pressure to cause slight flaring of the toes. The breadth was measured at the same spot as the width, but the calipers were rotated around the foot 90 degrees. Footpad size was calculated as width x breadth, and the degree of inflammation was expressed as the increase in size relative to the pre-infection measurement (day 0), obtained by the following formula: [(x-day 0)/day 0)], where x is the footpad size measurement for a given day post-infection.

### Statistical Analysis

SASS and GraphPad Prism Software (version 9.4.1) were used to analyze and graph data, with a p < 0.05 considered statistically significant. The change in joint inflammation was examined by treatment group, from baseline to the treatment period following CHIKV infection (baseline vs. days 16–19), using a random effects mixed model. This nests observations within subjects to account for correlated measurements on the same subjects over time. The ‘treatment’ time point was averaged across days 16–19. The treatment group-by-time interaction was also assessed to determine whether the slopes over time differed by group. The group and time main effects were tested to ascertain whether joint size differed across groups and time. Additionally, all analyses comparing groups were performed using a one-way ANOVA followed by Dunnett’s post-test comparing each treatment group to the control (PBS). Treg/Teff ratios were calculated, and mean ratios were analyzed by treatment group using a general linear model adjusted for sex. Histology composite scores were analyzed using a two-way ANOVA (mixed methods). Two independent ratings were done for each subject for the histology score, and test-retest reliability was assessed using the Pearson r, Bland-Altman plot and weighted kappa.

To determine sample size, raw data was used from Miner et al.’s 7-day change in foot-pad swelling in mice inoculated with CHIKV, comparing controls vs. mice treated with abatacept, to establish the expected effect size for IL-2 treatment on ankle swelling [15]. Their effect size was 0.97. In order to have a power > 0.95 to detect this effect in a 2-tailed (between-groups) t-test with an alpha of 0.05, our study needed at least 29 mice per treatment group.

## Data Availability

The datasets generated and analyzed during the current study are available from the corresponding author upon reasonable request.

## Acknowledgments

Biorender.com was used to create original figures. Research reported in this publication was supported by the National Institute of Arthritis and Musculoskeletal and Skin Diseases of the National Institutes of Health (Award Number K23AR076505) and the Pharmaceutical Research and Manufacturers of America (PhRMA) Foundation Research Starter Grant in Translational Medicine and Therapeutics. The content is solely the responsibility of the authors and does not necessarily represent the official views of the National Institutes of Health nor the PhRMA Foundation

## Author Contributions

SRT, AJP, GLS, CNM, and AYC were involved in the experimental design. SRT, AJP, PSL, and AYC performed experiments and acquired data. AMS interpreted histology slides and scored tarsal joints. SRT, AMP, and RA analyzed the data. SRT wrote the manuscript and prepared figures. All authors reviewed the manuscript.

## Additional Information

The authors declare no competing interests.

