## Supplemental Figure 1 for "Effects of IL2/anti-IL2 antibody complex on chikungunya virus-induced arthritis in a mouse model"

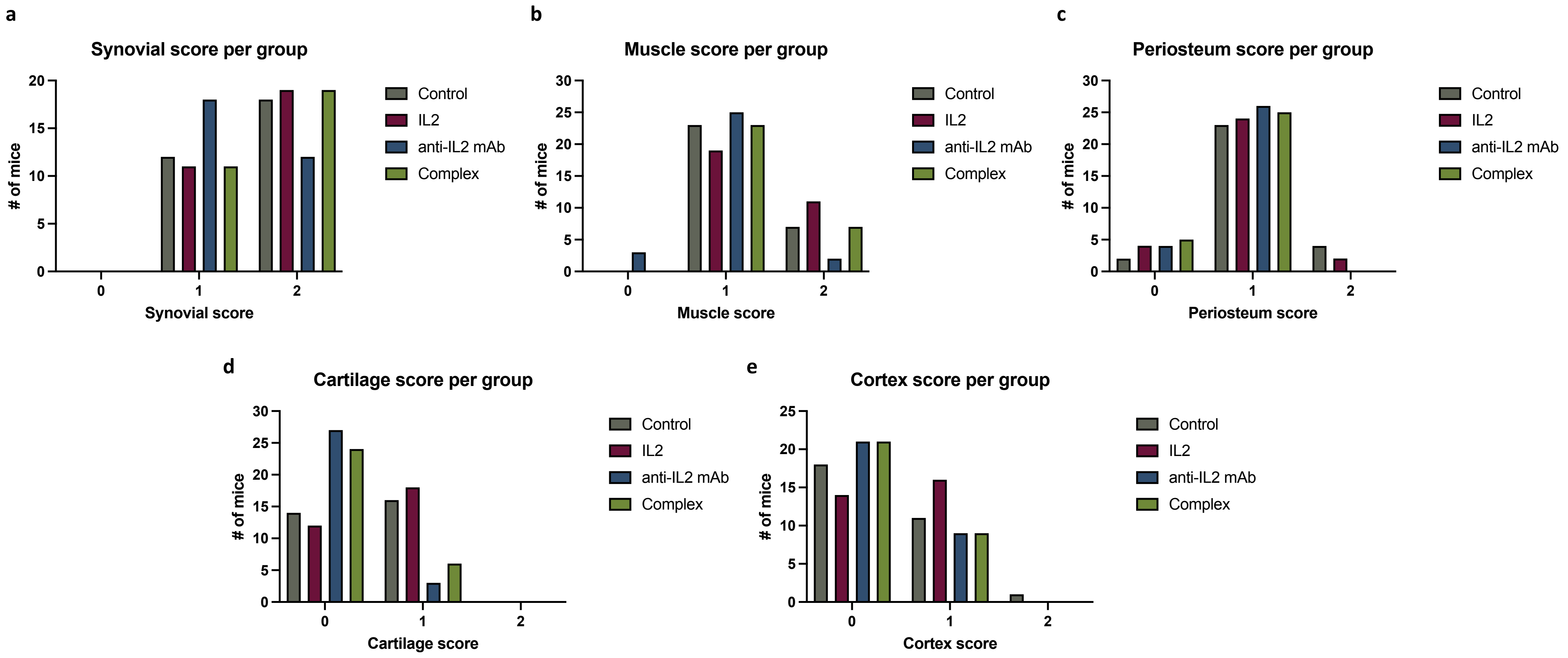


**Supplemental Figure 1.** Histological disease scores by component. Bars represent the number of mice per treatment group that received a score of 0, 1, or 2 for each histological component. Scores of 0 (no injury/inflammation) to 2 (significant injury/inflammation) were assessed for each histological component, including the synovium (A), skeletal muscle and soft tissue (B), periosteum (C), articular cartilage (D), and cortical bone (E).
